# Mixed effects modeling of the synergy between immune checkpoint and thrombin inhibitors in a preclinical mouse model of colon cancer

**DOI:** 10.64898/2026.07.29.741459

**Authors:** Nathanael Cox, Indrani Nayak, Zihai Li, Jayajit Das

## Abstract

Immune checkpoint blockade (ICB) therapy has revolutionized cancer treatment, though it is still effective in only about 20–40% of patients. To increase the efficacy of ICB therapy, combinations of ICB antibodies such as anti-PD1 and anticoagulants (e.g., thrombin inhibitors) have been studied in preclinical mouse models and clinical trials. We developed a Bliss-type analysis to quantify the synergy between anti-PD1 and dabigatran etexilate using published tumor growth data from a mouse model. We then developed a minimal mechanistic computational model to quantitatively study the synergy between anti-PD1 and the thrombin inhibitor dabigatran etexilate in a published study (Metelli et al.) of a preclinical mouse model of colon cancer. Our model included tumor cells, CD8+ T cells, and the pleiotropic cytokine TGFβ, whose production in platelets is influenced by thrombin, and described a potential mechanism of interplay among these components in the tumor microenvironment. We performed nonlinear mixed-effects modeling to capture mouse-to-mouse variation in tumor growth under different treatment conditions. The estimated parameter values pointed to several underlying mechanisms of synergy, including increased expansion of CD8+ T cells in the tumor microenvironment in the presence of anti-PD1 and dabigatran etexilate. These predictions can be further validated in future experiments. Thus, the combination of longitudinal tumor growth data, mechanistic population dynamics modeling, and nonlinear mixed-effects modeling can be used to study synergy between ICB and other drugs in controlling tumor growth.

## Introduction

Immunotherapeutic strategies that exploit the immune system to eliminate tumors have revolutionized cancer treatment^1^. Adaptive immune cells such as T cells are a major orchestrator of generating anti-tumor immune responses. A hallmark of cancer progression is the induction of T cells in an exhausted or dysfunctional state in the tumor microenvironment (TME) where the exhausted T cells have reduced ability to mount anti-tumor responses. In this exhausted state, T cells upregulate checkpoint co-receptor proteins such as PD1 and CTLA4 on the cell surface which then induce inhibitory signaling in T cells. Immune checkpoint blockage (ICB) therapies use antibodies such as anti-PD1/PDL1 which bind to the checkpoint receptors (e.g., PD1) or their cognate ligands (e,g., anti-PDL1) expressed on tumor cells which have been found to restore anti-tumor responses in exhausted T cells in preclinical mouse models and in the clinic. However, despite many striking examples of success, the ICB therapies are still effective on only a moderate percentage (~20-40%) of patients^1,2^. To increase the efficacy ICB therapies are being combined with additional drugs such as anticoagulants in clinical trials^3^.

Thrombin is an enzyme involved in blood clot formation^4^. Biomarkers for thrombin generation have been found to be elevated in cancer patients and are negatively associated with survival^4,5^. Thrombin can activate platelets, which help repair damaged blood vessels and promote wound healing, but it has also been shown to induce pro-tumor functions by facilitating the generation of active TGFβ^6^, a multifunctional cytokine involved in suppressing immune responses in the TME^7^. Thrombin inhibitors, such as dabigatran etexilate, have demonstrated synergistic effects in controlling tumor growth in preclinical mouse models when administered in conjunction with chemotherapy and ICB drugs^6,8^. The effects of dabigatran and other anticoagulant drugs as synergistic drugs in ICB therapies have been evaluated in recent clinical trials with mixed results^3^.

Development of immunotherapies with greater efficacy requires a mechanistic, systems level understanding of how the pro- and anti-tumor immune responses in the TME, involving diverse innate and adaptive immune cells such as macrophages, platelets, T cells, B cells, and NK cells, are affected by ICB or anticoagulant drugs. This is particularly challenging because the interactions between immune and tumor cells is shaped by crosstalk between immune cells which is dynamically regulated by multifunctional cytokines such as TGFβ. As a result, targeting a specific protein with a drug (e.g., PD1 by anti-PD1) can have non-intuitive effects on multiple pro- and anti-tumor immune responses. Mechanistic and dynamical computational models can help interrogate these complex interactions. When trained with data, these models can provide an in silico framework to evaluate the consequences of different drug combinations in controlling tumor growth and to explore a mechanistic basis of drug synergy. Mechanistic mathematical models based on ordinary and partial differential equations (ODEs and PDEs)^9^ that describe interactions between tumor cells, CD8+ T cells, anti-PD1/PDL1 drug, and cytokines have been developed for modeling responses to ICB therapy and to identify biomarkers^10^ and kinetic parameters^11^ associated with treatment success. Synergistic effects between ICB and cancer vaccine^12^ or immunostimulants (e.g., muIL-2) ^13^ or radiotherapy^14^ has been investigated using ODEs, PDEs, and discrete time modeling that describe interactions among tumor cells, therapeutic agents, and specific immune cell populations. However, the responses to the same ICB therapy or ICB combined with other treatments show substantial variability among individuals^15,16^ and even among genetically identical animals in preclinical mouse models^17^. The sources of this variability are not well understood and may include differences in genetic background, age, and comorbidities. In preclinical animal models, variability can arise from differences in the composition of immune cell populations or randomness in interaction^15^ between immune and tumor cells.

Developing mechanistic models that capture this variability can help identify common patterns in tumor-immune cell interactions within a patient or animal cohort as drugs are administered. Furthermore, such models may enable the development of digital twins that incorporate inter-individual variations. Recent studies have begun to incorporate variability in ICB responses in cancers such as head and neck squamous cell carcinoma^18^ and bladder cancer^19^ using nonlinear mixed-effects (NLME) modeling^20^. In addition, tumor growth in preclinical mouse models treated with combinations of radiotherapy, DNA damage response inhibitors^21^, and ICB has been modeled using NLME approaches to determine optimal dosing regimens.

We have developed an NLME model of tumor growth kinetics in a preclinical mouse model treated with ICB (anti PD-1) and anticoagulant (dabigatran) drugs to investigate potential mechanisms of synergy between these therapies. The model successfully describes animal-to-animal variation in tumor growth and identifies testable hypotheses for drug synergy. This approach can be extended to investigate mechanisms of drug synergy in tumor growth in the context of inter-animal or inter-patient variability.

## Results

### Quantification of synergy between anti-PD1 and thrombin inhibitor dabigatran etexilate in regulating tumor growth

We used published (Metelli et al.^6^) longitudinal changes in tumor size in preclinical MC38 mouse model when anti-PD1, or thrombin inhibitor (dabigatran etexilate), or the combination of anti-PD1 and dabigatran etexilate were administered to quantitatively evaluate any synergy present between thrombin inhibitor dabigatran and anti-PD1 in controlling tumor growth in the mouse model. In the experiments, MC38 colon carcinoma cells were injected subcutaneously in wild type C57BL/6 mice and the growth of the tumor was studied when the mice were treated with thrombin inhibitor dabigatran etexilate alone, or with anti-PD1 alone, or with a combination of thrombin inhibitor dabigatran and anti-PD1 or were left untreated (control). We analyzed the tumor growth data from day 7 to day 30 post injection of MC38 cell lines. The tumor growth data showed the tumor sizes decreased the most for the treatment with dabigatran+anti-PD1, whereas treatment with anti-PD1 or dabigatran alone decreased the tumor size compared to the control. We developed and implemented an analysis, similar to that of Bliss’s analysis^22,23^, to evaluate any presence of synergy between dabigatran etexilate and anti-PD1 for the tumor growth (see Methods). The analysis assumes that the tumor grows exponentially between the measured time intervals, e.g., day 7 (=t_1_) and day 11(=t_2_). We derived an inequality for the presence of any synergy between anti-PD1 and dabigatran when the drugs are administered simultaneously, in terms of the fold changes in the tumor size in a time interval (see Materials and Methods for details). The fold change (f_A_(t_1_, t_2_)) in the average tumor size for a treatment A (e.g., dabigatran alone) in the time interval t_1_ to t_2_ is computed from the tumor sizes C_A_(t_1_) and C_A_(t_2_), at times t_1_ and t_2_ as f_A_(t_1_, t_2_)=C_A_(t_2_)/C_A_(t_1_). If the fold changes in the average tumor size in the time interval t_1_ and t_2_ for treatment A (e.g., dabigatran etexilate alone), treatment B (e.g., anti-PD1 alone), the combination treatment A+B (e.g., anti-PD1+dabigatran etexilate), and in the isotype mice where no drugs are administered are given by, f_A_(t_1_, t_2_), f_B_(t_1_, t_2_), f_A+B_(t_1_, t_2_), and f_0_(t_1_, t_2_), respectively, we showed when anti-PD1 and dabigatran are synergistic, f_A+B_(t_1_, t_2_) < [f_A_(t_1_, t_2_)× f_B_(t_1_, t_2_)]/f_0_(t_1_, t_2_). The inequality implies that the effect of the combined treatment (A+B) is greater than that of sum of the parts (A or B) to limit tumor growth.

Application of the above analysis to the data obtained from Metelli et al.^6^ showed that Dabigatran and anti-PD1 synergize (Figure 1) at intermediate times (17 to 20 days) during tumor growth and the synergy decreases during early and becomes negligible during late tumor growth. Next, we set up a minimal mechanistic population dynamic model to describe potential mechanisms that may underlie the synergy between anti-PD1 and dabigatran in regulating tumor growth.

**Figure 1.**
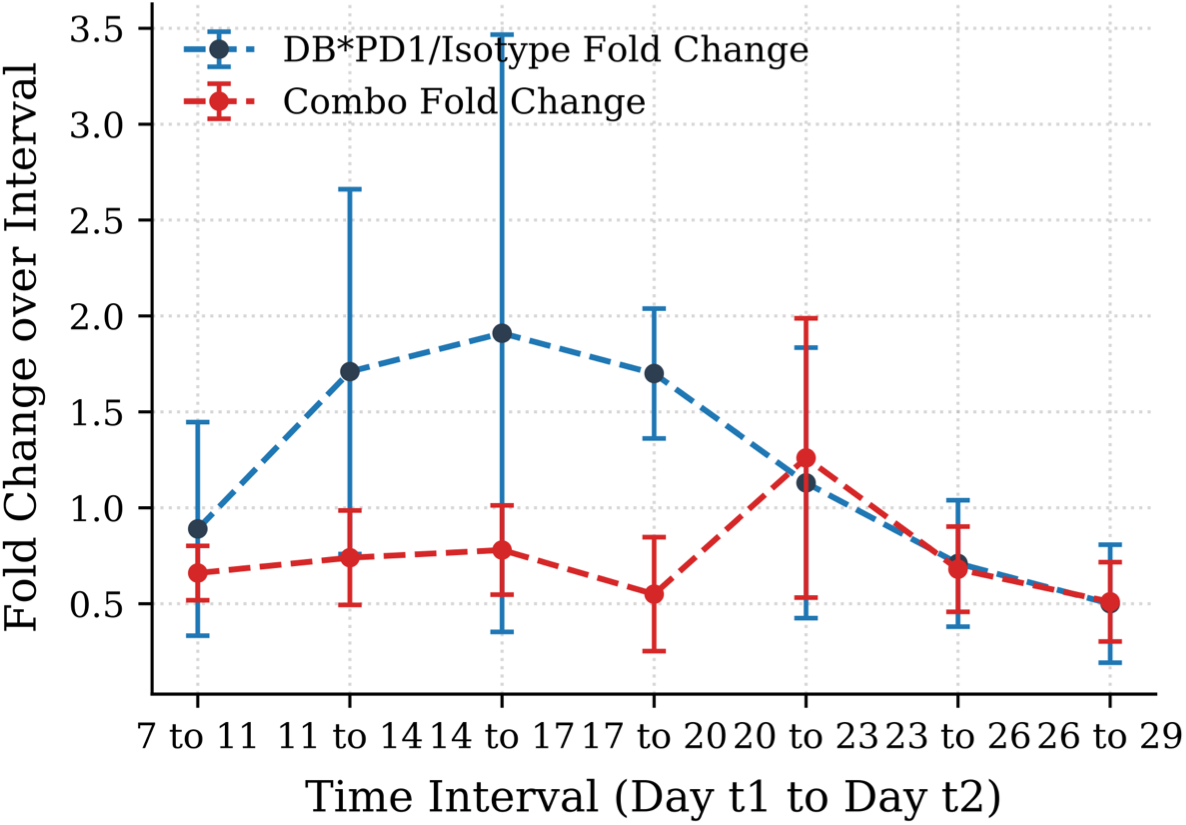
Quantification of the synergy between anti-PD1 and the thrombin inhibitor dabigatran in reducing tumor size. The scaled product of the fold changes, [*f*_DB_(*t*_1_, *t*_2_) × *f*_aPD1_(*t*_1_, *t*_2_)]/*f*_O_(*t*_1_, *t*_2_)(blue dashed line), is compared with the fold change *f*_DB+aPD1_(*t*_1_, *t*_2_)(red dashed line) in tumor area following combination treatment. Here, *f*_O_(*t*_1_, *t*_2_), *f*_DB_(*t*_1_, *t*_2_), *f*_aPD1_(*t*_1_, *t*_2_), and *f*_DB+aPD1_(*t*_1_, *t*_2_) denote the mean fold changes in tumor area between times *t*_1_and *t*_2_ across mice (n=9) for the isotype control (no treatment), dabigatran alone, anti-PD1 alone, and the combination of dabigatran and anti-PD1, respectively. Error bars represent the standard error of the mean.

### Mechanistic modeling of the interplay between anti-PD1 and thrombin inhibitor dabigatran etexilate in tumor growth

We developed a minimal population dynamic model using coupled ordinary differential equations to describe the time evolution of the populations of tumor and CD8+ T cells and the concentrations of cytokine TGFβ. The effects of thrombin inhibitor dabigatran and anti-PD1 antibodies are modeled implicitly. The interplay among these cell populations and TGFβ concentrations is incorporated into the model and is illustrated schematically in Figure 2A. In the model the tumor cells proliferate and are eliminated by tumor infiltrating cytotoxic responses of CD8+ T cells. The cytokine TGFβ is continuously produced in the tumor microenvironment^7^, and TGFβ binds to the TGFβ receptors expressed on CD8+ T cells^24^. The enzyme thrombin facilitates the release of active TGFβ from the complex formed by latent TGFβ and the cell-surface docking receptor GARP on platelets^6,25^. Dabigatran etexilate binds to the active site of thrombin^26^ and reduces the release of active TGFβ in the local environment^6^. Administration of dabigatran decreases the production rate of TGFβ, while administration of anti-PD-1 increases the rate of tumor lysis by cytotoxic CD8+ T cells. The kinetics of the cell populations and TGFβ1 concentration are described by coupled non-linear ODEs where we considered populations of tumor cells (*C*), tumor infiltrating CD8+ T cells (*T*), CD8+ T cells bound to TGFβ1 (*T*_*β*_), and TGFβ1 concentration (*β*). The effect of anti-PD1 is modeled implicitly. TGFβ1 is produced by platelets (implicit in the model) and bind to CD8+ T cells. In the model, the tumor cells proliferate with a fixed rate (ρ_C_) and the cancer cells are eliminated by the CD8+ T cells not bound or bound to TGFβ1 with rates 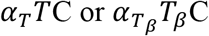, respectively. The addition of anti-PD1 increases the rates of tumor lysis by the CD8+ T cell populations *T* and *T*_*β*_ by *α*_*PD*_ *TC* and 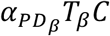, respectively, which describe the increased cytotoxicity of CD8+ T cells when the PD1 receptors are blocked by anti-PD1 molecules. The CD8+ T cells are stimulated by neoantigens derived from tumor cells and presented to T cells by antigen presenting cells in the tumor microenvironment and the draining lymph node which induces proliferation of the CD8+ T cells^27^. The neoantigens are derived from dead or lysed cancer cells^28^, therefore we reasoned that the stimulation resulting in T cell proliferation should be proportional to the cancer cell population *C*. In the model, the populations of CD8+ T cells unbound and bound to TGFβ, *T* and *T*_*β*_, respectively, proliferate with the rates *ρ*_*T*_*TC* and 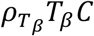. We do not consider the transport of T cells from the draining lymph nodes to the TME explicitly to keep the model simple. We considered the kinetics of the cytokine TGFβ1 in the TME and the draining lymph nodes. TGFβ1 is produced by diverse cell types including platelets and cancer cells. The production of TGFβ1 is represented by a constant production rate, *s*. The addition of dabigatran inhibits TGFβ1 production by platelets, which decreases the production rate *s* to (1-*k*) *s*, where 0≥*k*≥ 1. TGFβ1 can bind to TGFβ1 receptors on CD8+ T cells. The binding and unbinding of TGFβ1 are incorporated by second order mass action rate for binding, *k*_*on*_*Tβ* and a first order unbinding rate, *k*_*off*_ *T*_*β*_. TGFβ1 has a natural decay with a rate *λ*_*β*_*β*. A list of the parameters is provided in Table 1.

**Figure 2.**
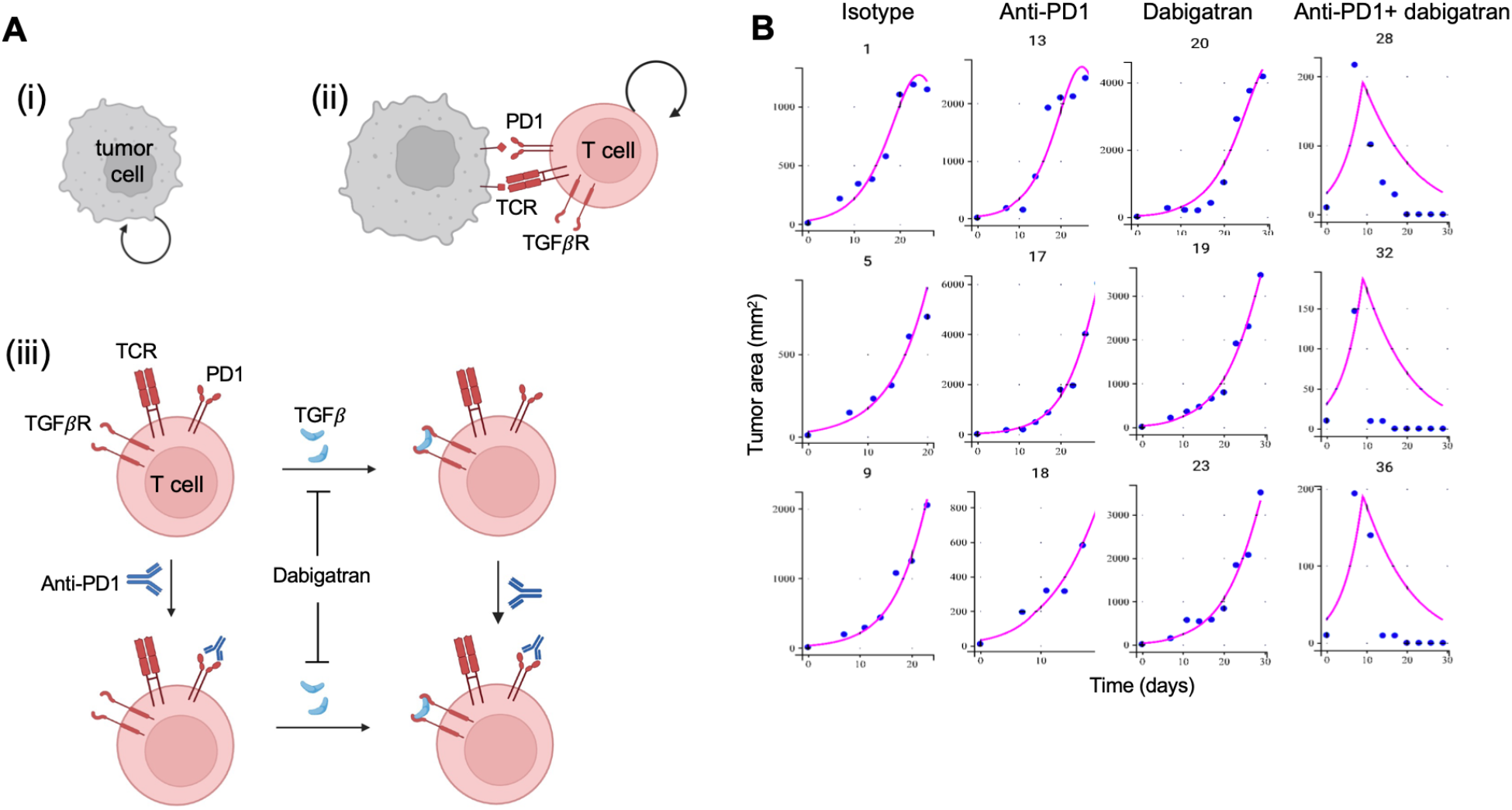
Modeling of tumor growth in the presence of anti-PD1 and the thrombin inhibitor dabigatran. (A) Schematic illustration of the interactions among T cells, tumor cells, anti-PD-1, and dabigatran. (i) Tumor cells proliferate within the tumor microenvironment (TME). (ii) T cells express TCR, PD1, and TGFβ receptors, interact with tumor cells, and undergo proliferation. (iii) PD1 and TGFβR expressed on T cells bind to extracellular anti-PD1 or TGFβ molecules. The presence of dabigatran etexilate reduces extracellular TGFβ concentration. Anti-PD1 blocks PD1 mediated inhibitory signaling, whereas dabigatran reduces TGFβ mediated immunosuppression. These drugs enhance T cell effector function, resulting in increased tumor cell lysis. (B) Model fits to tumor growth data from the MC38 mouse model under different treatment conditions. Each panel shows the tumor growth trajectory of an individual mouse.

**Table I:** List of model parameters and their values.

| Symbol | Parameter | Parameter Variation | Parameter Values [unit] |
| --- | --- | --- | --- |
| $\rho_C$ | Rate of tumor cell proliferation | Fixed | 0.35<br>[1/day] |
| $\rho_T$ | Proliferation rate of T cells not bound to TGFβ proliferation | Fixed | $1.88 \times 10^{-6}$<br>[1/(day×cell density)] |
| $\rho_{T\beta}$ | Proliferation rate of TGFβ bound T cells | Random | |
| $\alpha_T$ | Rate of tumor lysis by T cells not bound to TGFβ | Fixed | $6.8 \times 10^{-5}$<br>[1/(day×cell density<sup>2</sup>)] |
| $\alpha_{T\beta}$ | Rate of tumor lysis by T cells bound to TGFβ | Random | |
| $\alpha_{PD}$ | Rate of tumor lysis by T cells bound to anti-PD1 but not bound to TGFβ | Fixed | $10^{-5}$<br>[1/(day×cell density<sup>2</sup>)] |
| $\alpha_{PD\beta}$ | Rate of tumor lysis by T cells bound to anti-PD1 and bound to TGFβ | Random | |
| $k_{on}$ | Rate of TGFβ binding to T cells | Random | |
| $k_{off}$ | Rate of TGFβ unbinding from TGFβ bound T cells | Random | |
| $\lambda_{\beta}$ | Decay rate of TGFβ | Random | |
| $1 - k$ | Dabigatran-reduced TGFβ production rate | Random | |
| $s$ | Base production rate of TGFβ | Fixed | $60 \times 10^5$<br>[molecules/day] |
| $C_0$ | Initial concentration of Cancer cells | Fixed | $30 \times 10^5$<br>[# of cells] |
| $T_0$ | Initial concentration of T cells | Fixed | $3 \times 10^5$<br>[# of cells] |

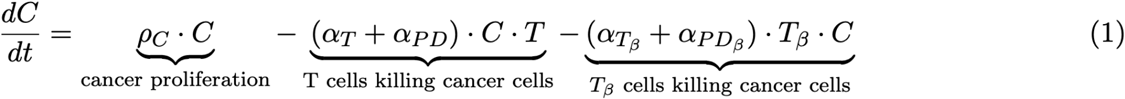

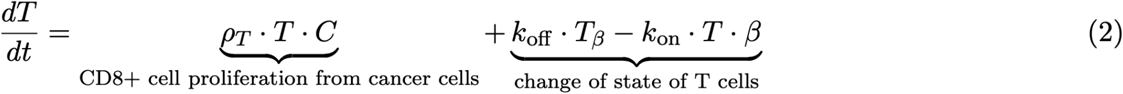

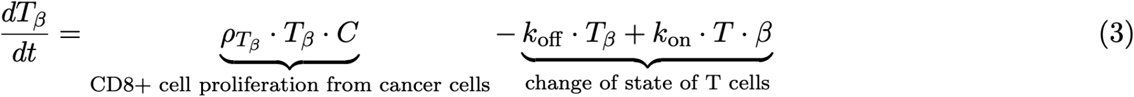

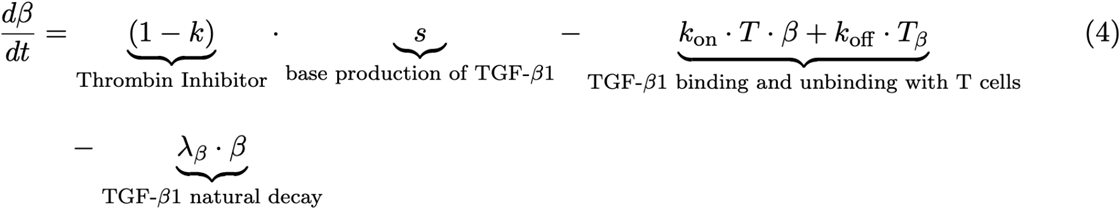

Next, we applied the above model (Eqs. 1-4) to describe the tumor growth kinetics reported by Metelli et al.^6^ (Figure 2B) under different treatment conditions. Eqs. (1-4) represent the combination treatment, in which both dabigatran and anti-PD1 are administered. To model isotype control, we set *α*_*PD*_, 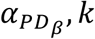 to zero in Eqs. (1) and (4). For anti-PD1 treatment alone, we set *k* to zero in Eq. (4), whereas for dabigatran treatment alone, we set *α*_*PD*_, 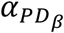 to zero in Eq. (1). The tumor growth kinetics in the mice showed large variations in particular when anti-PD1 and/or dabigatran were added in the treatment. Therefore, we developed a nonlinear mixed effects model which is capable of capturing the mouse-mouse variations of the tumor growth in the experiments.

### Nonlinear mixed effects modeling of MC38 tumor growth in mice

We developed a non-linear mixed effects (NLME) model using the ODEs in Eqs. (1-4) to describe the tumor growth kinetics in the untreated mice and the mice treated with dabigatran alone, or anti-PD1 alone, or with the combination dabigatran+anti-PD1. The NLME model contains population or fixed parameters (***θ***_pop_; a parameter vector is shown with a bold face symbol) which do not vary with individual mice and are held fixed, and random parameters, ***θ***_*i*_, which vary from mouse to mouse where each mouse is indexed by *i*. We considered ***θ***_pop_ ≡ {*T*_0_, *ρ*_C_, *ρ*_T_, *α*_T_, *α*_PD_, *s*} to be the fixed parameters, and 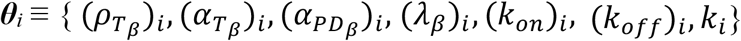 to be the random parameters (individual) that vary across mice. The random parameters were assumed to follow a log-normal distribution with means (**μ**) and standard deviations (**ω**). The number of tumor cells, (C_(data)_(t)), was estimated from the experimentally measured tumor area, (C_(expt)_(t)), using the approach described in ref. ^29^ for the MC38 mouse model. The residual differences between the estimated number of tumor cells, C_(data)_(t), and the ODE model predictions, C_(model)_(t) were assumed to be normally distributed with mean zero and variance *σ*^2^ at each observation time *t*. The likelihood function, *L*(*C*_(*expt*)_ (*t*), *θ*_*pop*_, *θ*_*i*_; *q*), where, *q* = {*σ*, ***μ, ω***}, was maximized to estimate the parameters. The fixed parameters were chosen to be of the same order of magnitude of the parameters used in our experimentally validated population dynamic model developed for tumor growth in the preclinical MC38 mouse model^29^. Software package Monolix was used for the parameter estimation. Further details regarding the parameter estimation are provided in the Materials and Methods section.

The model with fixed and randomly varying parameters across the animals describes the animal-animal variations of the tumor growth in four different conditions: isotype, with dabigatran, with anti-PD1, and with anti-PD1+dabigatran. The average values of tumor growth measurements show increase of tumor size over three weeks in isotype mice, and the growth slows down when dabigatran or anti-PD1 is administered. However, when anti-PD1+dabigatran is administered the average tumor size starts decreasing after about 10 days. The tumor growth in individual animals shows large variations, and the mixed effects model is able to capture these variances (Figure 2B and Supplementary Figure S1). The R^2^ value at 0.53 shows the excellent agreement with the model and the data (Supplementary Text 1, Supplementary Figure S2). Next, we analyzed the variations of the random parameters for each case to determine processes that are associated with the synergy between anti-PD1 and dabigatran. The distributions and pair-correlations between the random parameters are shown in the Supplementary Figure S3. The mean increase in the tumor cell killing rate of TGFβ-bound CD8+ T cells in the presence of anti-PD1 (i.e., CD8+ T cells bound to both TGFβ and anti-PD1 or CD8+ T cells bound to TGFβ but not bound to anti-PD1) does not show any statistically significant difference when anti-PD1 is added alone or with dabigatran (Figures 3A and 3B). Similarly, the average rate of production of TGFβ (or 1-*k*) does not display any statistically significant difference between the anti-PD1 and anti-PD1+dabigatran cohorts (Figure 3C). However, the rate of proliferation of TGFβ bound CD8+ T cells showed a statistically significant increase in the anti-PD1+dabigatran cohort, compared to the anti-PD1 alone, dabigatran alone, or the isotype cohort (Figure 3D). The average value of the random rate parameters associated with binding and unbinding of TGFβ with TGFβ receptors, and the decay rate of TGFβ, do not show any statistically significant differences in the four above cohorts (Supplementary Figure S4). Thus, the estimated parameters indicate that the presence of anti-PD1 and dabigatran increases the proliferation rate of TGFβ bound CD8+ T cells.

**Figure 3.**
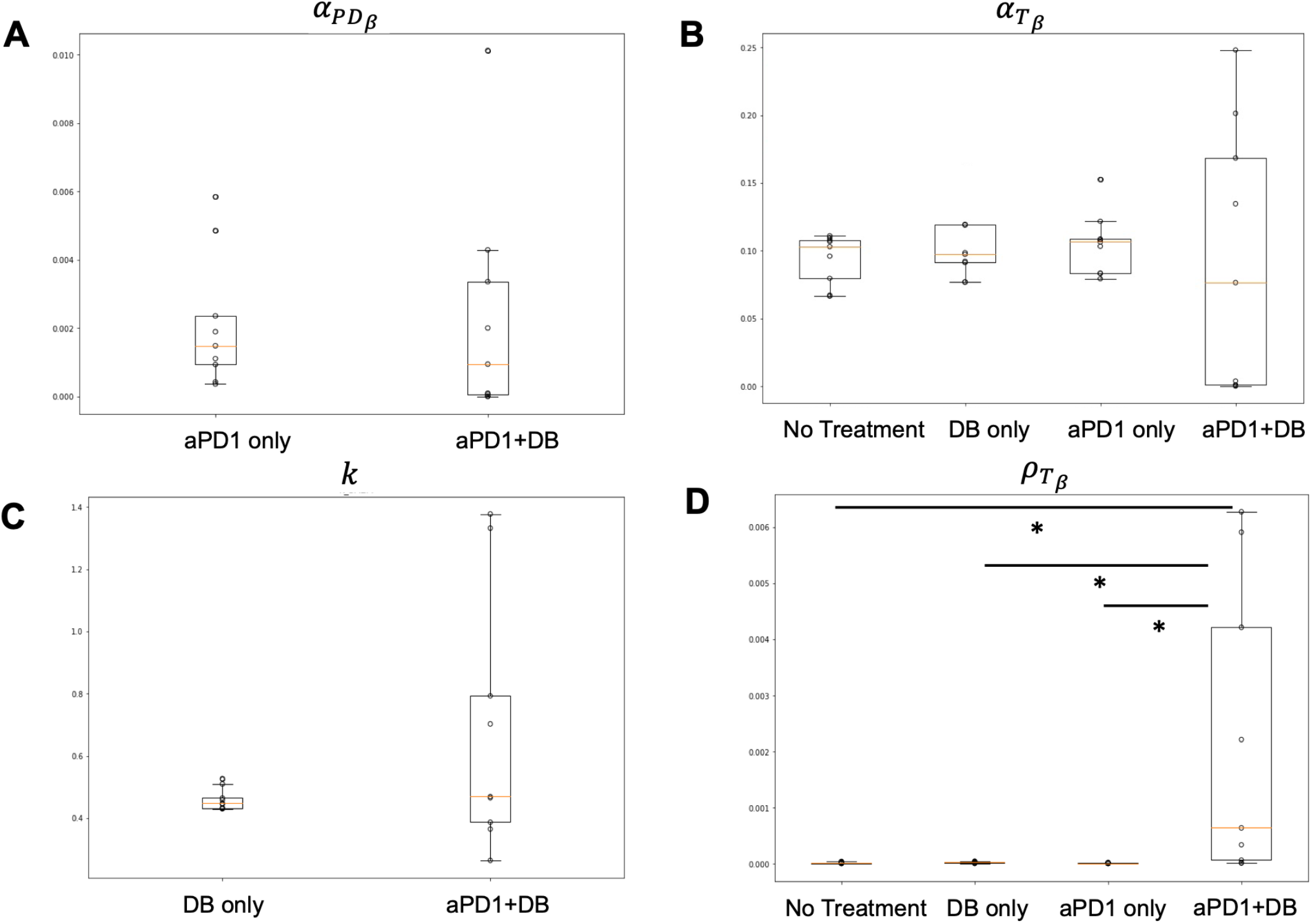
Parameter estimation in the NMLE model. (A) Shows the values of random parameter 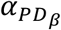, the increase in the rate of lysis of the tumor cells by TGFβ bound CD8+ T cells with anti-PD1, for anti-PD1 only or the combination (dabigatran etexilate+anti-PD1) case. (B) Shows the values of random parameter 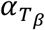, the rate of lysis of tumor cells by TGFβ bound CD8+ T cells, for the isotype (no treatment), dabigatran etexilate only, anti-PD1 only, or the combination (dabigatran etexilate+anti-PD1) case. Though individual values for the combination can be larger than all the other cases, the mean values are not separated by statistical significance. (C) Shows the values of random parameter *k*, the change in the TGFβ or dabigatran etexilate only or the combination (dabigatran etexilate+anti-PD1) case. (D) Shows the values of random parameter 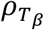, the rate of proliferation of the TGFβ bound CD8+ T cells, for the isotype (no treatment), dabigatran etexilate only, anti-PD1 only, or the combination (dabigatran etexilate+anti-PD1) case. The mean value for the combination is larger than all the other cases. The asterisk (*) denotes a statistically significant difference (P<0.05).

## Discussion

We developed a minimal model to capture the synergy between anti-PD1 ICB and an anticoagulant in controlling tumor growth in a preclinical mouse model. The model successfully described the drug synergy as well as captured the mouse-to-mouse variations of the tumor growth. The model describes interaction between tumor and CD8+ T cells in the TME minimally, thus approximates several detailed interactions such as neoantigen stimulated T cell activation via the rate parameters associated with the population dynamic model. Similar approximations have been made in previously developed Lotka-Volterra type models for describing tumor-immune interactions^11,29-31^. The presence of anti-PD1 blocks the inhibitory interaction between inhibitory ligands PD-L1 expressed by the tumor cells or other immune cells (e.g., macrophages) and thus suppresses production of ‘exhausted’ or dysfunctional CD8+ T cells which show a reduced anti-tumor response^32^. The model minimally describes this effect by increasing the rate of tumor cell lysis by CD8+ T cells in the presence of anti-PD1. The synergy between anti-PD1 and the anticoagulant dabigatran etexilate is captured by modeling the effect of the cytokine TGFβ whose production is suppressed by the presence of dabigatran etexilate.

The variations in the tumor growth across mice are generated by random parameters that change from mouse-to-mouse. In our model we assigned several parameters such as the proliferation rate of TGFβ bound CD8+ T cells, or the rate of tumor lysis by TGFβ bound CD8+ T cells as random parameters. The choice of the random parameters was based on selection of random and fixed parameters that produced a convergence in parameter estimation. Since we fitted the population of tumor cells in our scheme and did not fit the T cell populations as the data were not available, there was many instances where a choice of fixed and random parameters would lead to unbound variations of the random parameters; these cases were not considered in our analysis.

The estimated random parameters showed statistically significant (p<0.05) increase in the population mean of the TGFβ bound CD8+ T cells when both anti-PD1 and dabigatran etexilate are present compared to the other cases (anti-PD1 alone, dabigatran etexilate alone, or isotype). The changes in the population mean of the other parameters such as rate of lysis by TGFβ bound CD8+ T cells in the presence of dabigatran alone, or anti-PD1 alone or the combination dabigatran+anti-PD1 do not show any statistically significant difference. Metelli et al.^6^ found dabigatran increased T cell infiltration in the TME in the MC38 model. Another study using a melanoma bearing mouse model showed increased infiltration of CD8+ T cells in the TME when anti-PD1 and an anticoagulant (STP3725) is combined^33^. The increase in the rate of proliferation of TGFβ T cells which increases the size of the T cell population in the TME in the model could point to such an effect mechanistically. Similarly, the absence of any further increase the rate of tumor lysis when dabigatran etexilate is added to anti-PD1 in the model leads to a testable prediction that dabigatran has minor effect in increasing the cytotoxicity of anti-PD1 bound CD8+ T cells. Thus, our approach can be utilized to develop mechanistic in silico models that can describe tumor growth in preclinical mouse models in the presence of synergizing drug combinations and can make testable predictions.

## Materials and Methods

### Quantification of drug synergy in tumor growth

We consider a pure birth model of tumor growth where the MC38 tumor cells are dividing in the wild type mice (control), in presence of anti-PD1 (case A) or dabigatran (case B) or both (case A+B). In the pure birth model, tumor cells starting with an initial population of n_0_ at t=0, divide with a rate k, i.e., a tumor cell in the population of n number of tumor cells divide with a rate k to create a population of n+1 tumor cells or n ***→*** n + 1. Since the cell division process is stochastic in nature, to have n number of tumor cells at a time t starting from n_0_ cells at t=0 occurs with a conditional probability, P(n,t|n_0_,0) which we denote as P(n,t) to simplify the notation. The time evolution of P(n,t) is described by a set of coupled linear ordinary differential equations, also known as the Master equation, as follows.

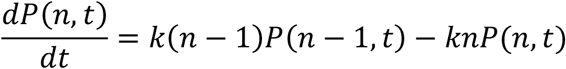

which can be solved exactly, and the solution is given by,

*P*(*n, t*) = *n*_O_*e*^−*kt*^(1 − *e*^−*kt*^)^*n*–1^, and the average number of tumor cells at any time t, is given by, 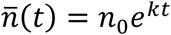. Therefore, the fold change *f*(t_2_, t_1_) in tumor size for a time interval *t*_2_ to 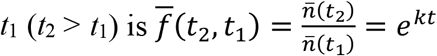. Since the experiments show mouse to mouse variation of the tumor growth, we compute the average of 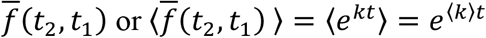 across animals in the cohort where ⟨… ⟩ denotes the average over animals. To keep the notation simple, we use *f*(*t*_2_, *t*_1_) = *e*^*kt*^ to represent 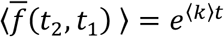.

If the rates for tumor growth are *k*_0_, *k*_0_-*k*_A_, *k*_0_-*k*_B_, and, *k*_0_-*k*_A+B_ for the control, case A, case B, and case A+B, respectively, then the fold changes in the tumor populations for a time interval T=(*t*_2_-*t*_1_) these cases are given by, 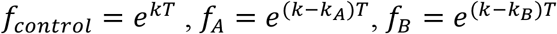, and, 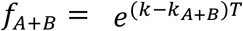, respectively. In the absence of any synergy between case A and case B, *k*_*A*+*B*_ = *k*_*A*_ + *k*_*B*_, therefore, *f*_*A*+*B*_ × *f*_*control*_ = *f*_*A*_*f*_*B*_. When synergy between the case A and B are present, *k*_*A*+*B*_ > *k*_*A*_ + *k*_*B*_, thus, *f*_*A*+*B*_ × *f*_*control*_ < *f*_*A*_*f*_*B*_.

### Nonlinear mixed effects modeling

The NLME calculations were performed using the software package Monolix (https://www.simulations-plus.com/software/monolix/monolix/). The random parameters were assumed to follow log-normal distributions with mean **μ** and variance **ω**, which were estimated. Tumor cell population measurements obtained from individual mice at days 7, 11, 14, 17, 20, 23, 26, and 29 were used as the observed data for model calibration. Model parameters were estimated in Monolix using the Stochastic Approximation Expectation Maximization (SAEM) algorithm for maximum likelihood estimation (MLE). The ODEs were solved numerically using a 4^th^ order Runge-Kuta method. The tumor and T cell counts in the ODEs were divided by a factor of 10^5^ to avoid large numbers during the numerical solution of the ODEs.

### Estimation of the tumor cell population from experiments

We estimated the population of the tumor cells using the measured tumor area in Metelli et al.^6^ and the flow cytometry measurements for the MC38 tumor growth mouse model as described in our previous work^29^. Briefly, the tumor area is converted into tumor volume following an estimation procedure in ^29^, and flow cytometry experiments show that roughly 70% of the tumor volume is occupied by the tumor cells which was used to estimate the tumor populations.

## CRediT authorship contribution statement

**Nathanael Cox**: Writing – review & editing, Writing – original draft, Methodology, Investigation, Visualization, Validation, Formal analysis, Data curation, Software, Resources, Conceptualization; **Indrani Nayak**: Methodology and Writing-review. **Jayajit Das**: Writing-review & editing, Writing – original draft, Visualization, Supervision, Project administration, Methodology, Funding acquisition, Conceptualization.

## Acknowledgements

The work was funded by the Research Institute at the Nationwide Children’s Hospital. We thank Darren Wethington for discussions. ChatGPT (OpenAI) was used occasionally for language editing for improving readability.

