## Supplemental Figures for "Mixed effects modeling of the synergy between immune checkpoint and thrombin inhibitors in a preclinical mouse model of colon cancer"

### Evaluation of Model Performance Following Outlier Exclusion

Including all data points resulted in a poor coefficient of determination ($R^2$=-43), driven by a small number of extreme outliers. Applying increasingly stringent outlier thresholds improved the agreement between predicted and observed values. By excluding predicted values that were greater than 15,000, 10,000, and 5,000, the corresponding $R^2$ values were 0.529, 0.696, and 0.702, which excluded 3, 5, and 8 data points, respectively (Supplementary Figure S2).

Because mouse 33 exhibited unusually large deviations in tumor area over time (Supplementary Figure S2E), exclusion of its three outlier measurements was considered justified. This resulted in a substantial improvement in model performance, with the coefficient of determination increasing to $R^2$ = 0.529 (Supplementary Figure S2B).

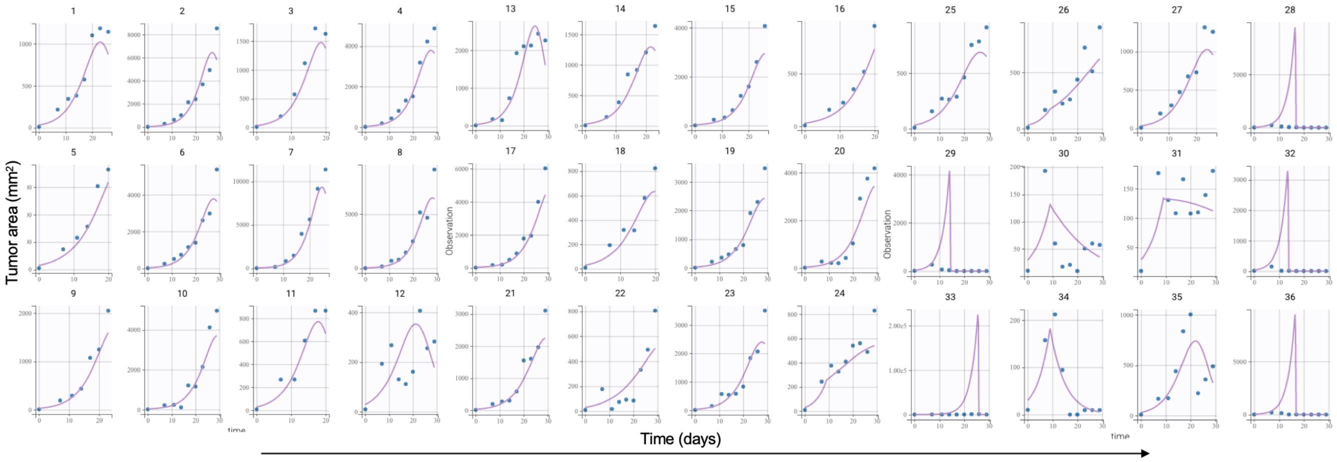


**Figure S1.** Model fits to tumor growth data from the MB38 mouse model. Each panel shows the tumor growth trajectory of an individual mouse. Images 1-9 are mice that received no treatment. Images 10-18 are mice that received on Dabigatran treatment. Images 19-27 are for mice that received only PD-1 inhibitor. Images 28-36 are the mice that received both.

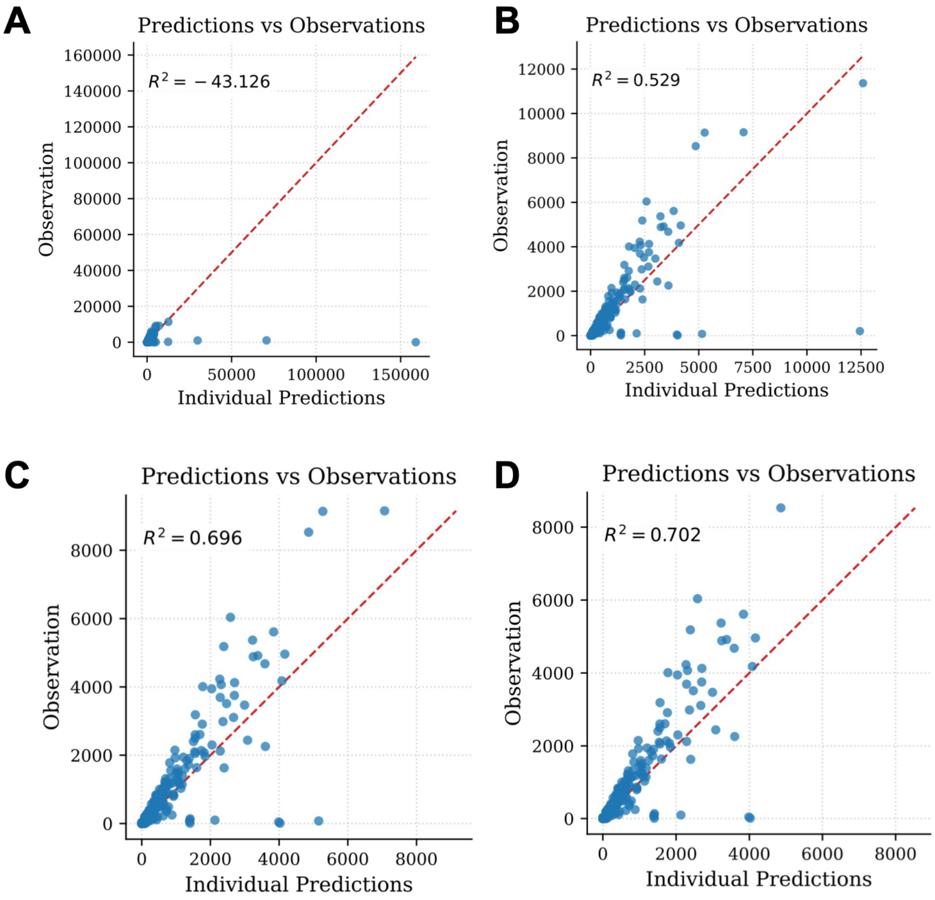


**Figure S2. Evaluation of model performance following outlier exclusion.** Coefficient of determination ($R^2$) values are shown for observed versus individual predicted tumor areas after excluding outliers. (A) All data points included (all mice, no outliers removed). (B) After removing 3 outliers (all three data points are from mouse 33 in the anti-PD1 + dabigatran combination treatment group). (C) After removing 5 outliers (data points are from mouse 7 in the untreated group and mouse 33 in the anti-PD1 + dabigatran combination treatment group). (D) After removing 8 outliers (data points are from mice 7 and 8 in the untreated group and mouse 33 in the anti-PD1 + dabigatran combination treatment group).

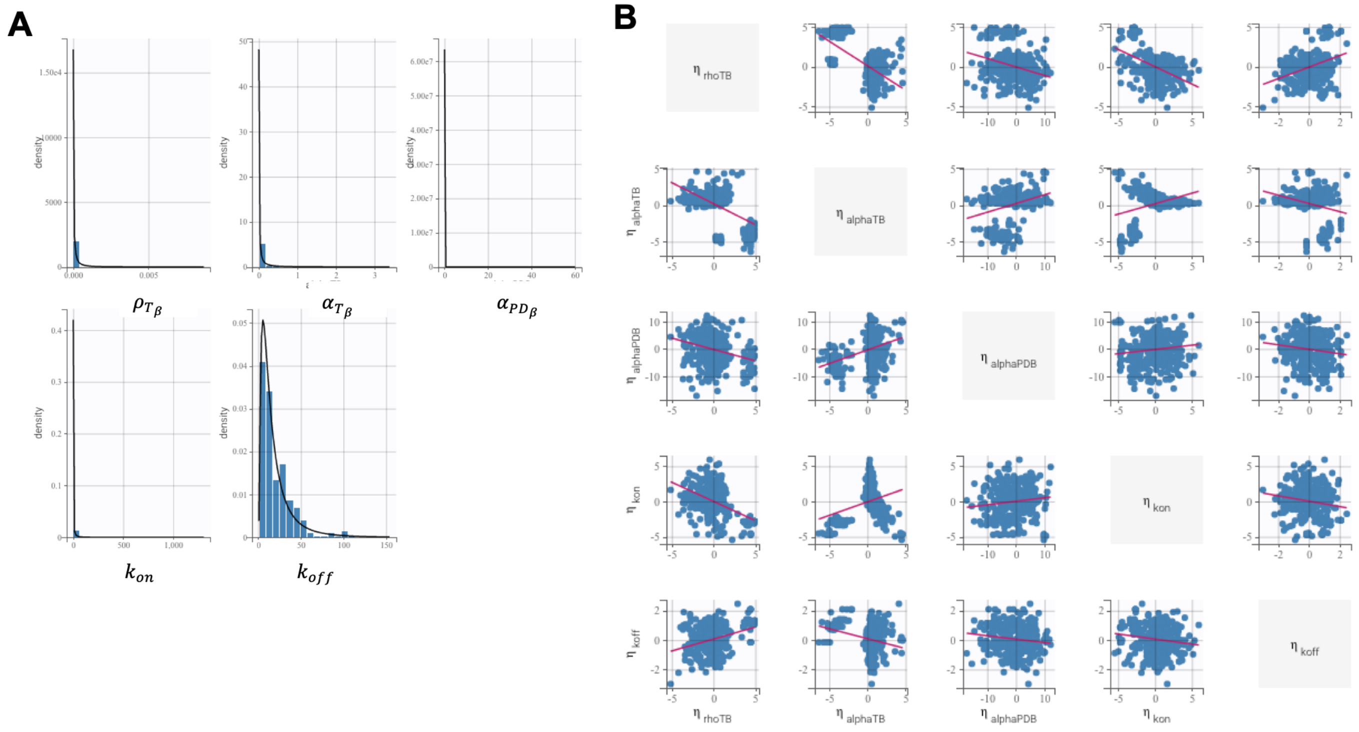


**Figure S3.** (A) Probability distribution of estimated parameters in the NMLE model. (B) Here each axis represents the deviation of each parameter for an individual mouse from the population parameter value. This compares the deviations in each parameter with the deviations in each other parameter for the mice.

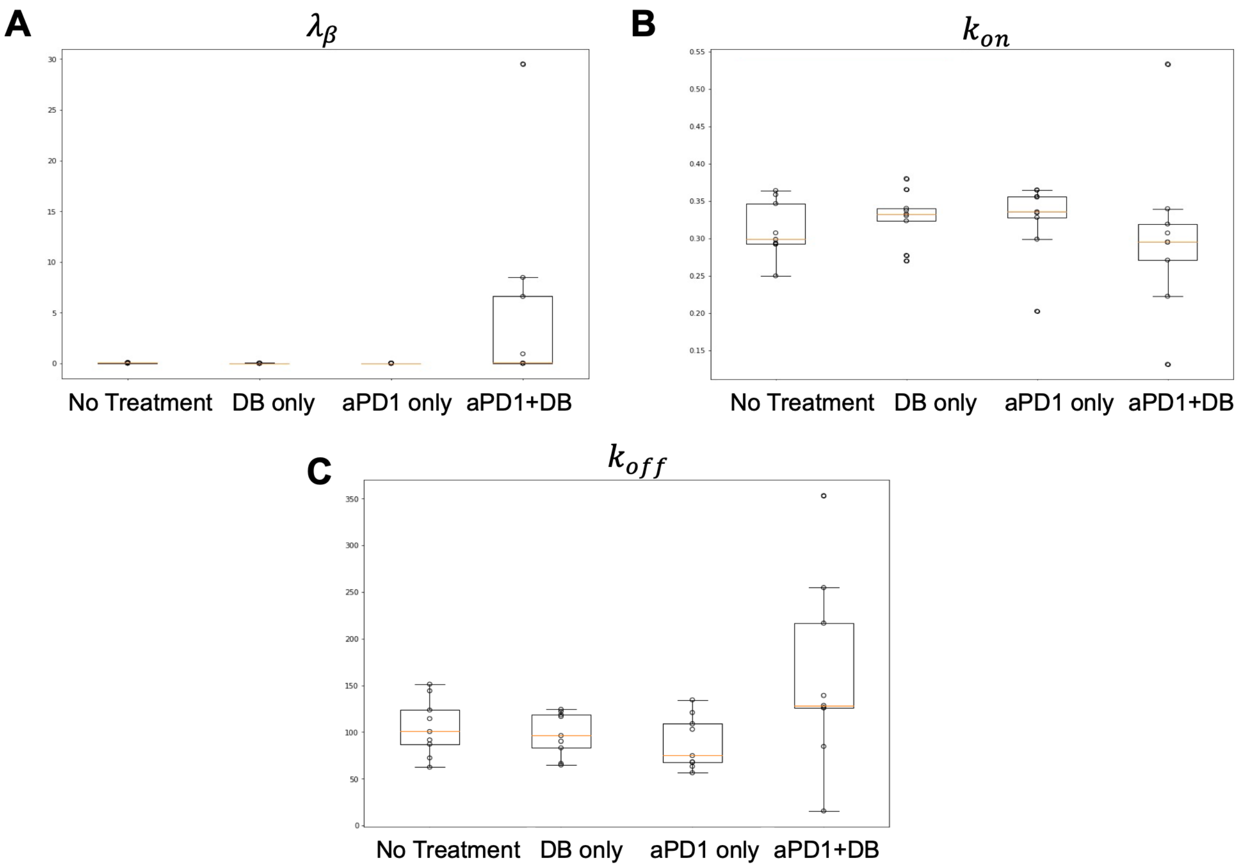


**Figure S4. Parameter estimation in the NMLE model.** (A) Shows the values of random parameter $\lambda_\beta$ , the decay rate of TGFβ for the isotype (no treatment), dabigatran etexilate only, anti-PD1 only, or the combination (dabigatran etexilate+anti-PD1) case. The mean value for the combination is larger than all the other cases. (B) Shows the values of random parameter $k_{on}$, the rate of TGFβ binding to CD8+ T cells with anti-PD1, for anti-PD1 only or the combination (dabigatran etexilate+anti-PD1) case. (C) Shows the values of random parameter $k_{off}$, the rate of TGFβ unbinding from TGFβ bound T cells in the TGFβ or dabigatran etexilate only or the combination (dabigatran etexilate+anti-PD1) case.
